# Ongoing evolution of the *Mycobacterium tuberculosis* lactate dehydrogenase reveals the pleiotropic effects of bacterial adaption to host pressure

**DOI:** 10.1101/2023.10.09.561592

**Authors:** Sydney Stanley, Xin Wang, Qingyun Liu, Young Yon Kwon, Abigail M Frey, Nathan D Hicks, Andrew J Vickers, Sheng Hui, Sarah M Fortune

## Abstract

The bacterial determinants that facilitate *Mycobacterium tuberculosis* (Mtb) adaptation to the human host environment are poorly characterized. We have sought to decipher the pressures facing the bacterium *in vivo* by assessing Mtb genes that are under positive selection in clinical isolates. One of the strongest targets of selection in the Mtb genome is *lldD2*, which encodes a quinone-dependent L-lactate dehydrogenase (LldD2) that catalyzes the oxidation of lactate to pyruvate. Lactate accumulation is a salient feature of the intracellular environment during infection and *lldD2* is essential for Mtb growth in macrophages. We determined the extent of *lldD2* variation across a set of global clinical isolates and defined how prevalent mutations modulates Mtb fitness. We show the stepwise nature of *lldD2* evolution that occurs as a result of ongoing *lldD2* selection in the background of ancestral lineage defining mutations and demonstrate that the genetic evolution of *lldD2* additively augments Mtb growth in lactate. Using quinone-dependent antibiotic susceptibility as a functional reporter, we also find that the evolved *lldD2* mutations functionally increase the quinone-dependent activity of LldD2. Using ^13^C-lactate metabolic flux tracing, we find that *lldD2* is necessary for robust incorporation of lactate into central carbon metabolism. In the absence of *lldD2*, label preferentially accumulates in methylglyoxal precursors dihydroxyacetone phosphate (DHAP) and glyceraldehyde-3-phosphate (G3P) and is associated with a discernible growth defect, providing experimental evidence for accumulated lactate toxicity via a methylglyoxal pathway that has been proposed previously. The evolved *lldD2* variants increase lactate incorporation to pyruvate but also alter flux in the methylglyoxal pathway, suggesting both an anaplerotic and detoxification benefit to *lldD2* evolution. We further show that the mycobacterial cell is transcriptionally sensitive to the changes associated with altered *lldD2* activity which affect the expression of genes involved in cell wall lipid metabolism and the ESX-1 virulence system. Together, these data illustrate a multifunctional role of LldD2 that provide context for the selective advantage of *lldD2* mutations in adapting to host stress.

## Introduction

*Mycobacterium tuberculosis* (Mtb) is currently the leading infectious cause of death globally (1). The temporal and geographic extent of the tuberculosis (TB) pandemic reflects remarkable adaption to human populations (2,3). Mtb is distinguished by limited genetic diversity due to a low mutation rate and minimal horizontal gene transfer and recombination events (4-6). However, the 7 genetically divergent human-adapted lineages of the *Mycobacterium tuberculosis* complex (MTBC) have distinct epidemiologic characteristics (7), suggesting that the mutations that define these lineages and sub-lineages are functionally important. Mtb strains also continue to acquire mutations that may provide selectively advantageous in the face of host environment and/or antibiotic pressures (8-11). Studies defining the phenotypic consequences of positive selection as a result of drug pressure have uncovered altered antibiotic sensitivity phenotypes conferring resistance, tolerance, and resilience (8,12-19). Considerably less is known about the functional consequence of positive selection driven by host pressure or the impact of lineage and sub-lineage defining mutations on host fitness.

In the current study, we sought to delineate the impact of functional genetic diversity in response to host challenge. We focus here on *lldD2* (*rv1872c*), a Mtb gene that encodes the lactate dehydrogenase (LDH) LldD2, which has been shown to be required for Mtb grown on lactate as a sole carbon source (20). Previous studies have shown that *lldD2* has one of the strongest signatures of nonantibiotic-related diversifying selection and mutations are prevalent in clinical strains (8,10, 21). There are lineage and sub-lineage defining mutations in *lldD2* which were acquired thousands of years ago (10), and mutations which continue to emerge contemporaneously in clinical strains from different branches of the phylogenetic tree (8,10,21). In addition, *lldD2* is under positive selection in *M. kansasii*, an environmental nontuberculous mycobacterium with an increasing incidence of human infection, which may suggest that *lldD2* is important for mycobacterial adaption to the human host (22).

Lactate is a prominent feature of the host environment; it is a significant contributor to tricarboxylic acid cycle (TCA cycle) metabolism across several tissue types, especially the lung (23). Further, lactate can be considered an immune-regulated host feature. Mtb infection in mice and IFNγ activated macrophages induce the Warburg effect, in which host lactate is produced as a result of aerobic glycolysis (24,25). Furthermore, lactate accumulation has been identified in mouse and guinea pig lung granulomas (26,27). Deletion of *lldD2* impairs Mtb survival in macrophages, demonstrating a requirement for lactate metabolism *in vivo* (20). Consistent with the *in vivo* relevance, a *lldD2* promoter mutation that is prevalent among clinical Mtb strains is associated with increased transmissibility (28).

It is unclear why *lldD2* is necessary for Mtb infection. In this study, we assess the functional effects of *lldD2* evolution. Utilizing a panel of Mtb clinical isolates and isogenic strains, we demonstrate that selected *lldD2* variants augment Mtb growth in lactate, modulate metabolic flux, and have striking secondary effects on expression of critical virulence systems. Taken together, these data inform our understanding of the evolution of Mtb to the human host and critical processes in Mtb pathogenesis.

## Results

### Stepwise evolution of *lldD2* increases Mtb fitness in lactate

An early study analyzed genome sequences from 220 MTBC isolates and characterized multiple genotypes of *lldD2* (10). They identified that three *lldD2* codons (3, 109, and 253) were subject to diversifying selection, and the V253M mutation in particular had been acquired by the ancestor of L2 before lineage diversification occurred (10). Subsequently, our work and that of others, which involved much larger datasets of MTBC whole-genome sequences, have revealed that *lldD2* is under positive selection, and this selection process is still ongoing in contemporary MTBC strains (8,21). Together, these observations suggest that selective pressure on *lldD2* has persisted since the very beginning of MTBC diversification, enduring over thousands of years.

To gain deeper insights into the evolutionary dynamics of the *lldD2* gene within the MTBC population, we assembled a comprehensive collection of 296 MTBC strains, which collectively represented the major lineages and sub-lineages (29). Employing the *lldD2* sequence derived from the inferred most recent common ancestor of the MTBC, we reconstructed the gene’s evolutionary history within the MTBC population. We find that there are root mutations in *lldD2* in every Mtb lineage except L1 and L6. The modern Mtb lineages (L2, L3, and L4) and L7 share an ancestral A176V mutation and L4 shares an additional lineage-defining A59G mutation (Fig. 1). L5 strains, which are highly prevalent in West Africa, derived an A2A and A237S from their common ancestor (Fig. 1). Notably, we also observed a remarkable pattern of stepwise evolution in *lldD2*, resulting in the accumulation of up to four distinct mutations in some sub-lineages or clades (Fig. 1). For instance, following the acquisition of the A176V and A59G mutations, a specific clade within L4 further accumulated V253M and L96F mutations (Fig. 1). Derived mutations span the *lldD2* promoter and coding regions but the V3I, V253M, and the -18 G>T promoter mutations in particular have recurrently evolved independently among MTBC strains (Fig. 1, Fig. 2A) (10,21,28). In an analysis of over 50,000 global MTBC clinical strains encompassing lineages L1-L7, these three mutations account for 52% of the 17,000 *lldD2* homoplastic or clade-defining mutations occurring across 216 different loci (Fig. 2B). The convergent signals of these homoplastic mutations strongly imply their adaptive role during the evolution of MTBC. The stepwise accumulation of multiple adaptive mutations suggests the existence of a complex and dynamic fitness landscape governing the evolution of *lldD2*. In this landscape, a single ancestral lineage-defining mutation may enhance fitness, while specific additional mutations can drive fitness even higher, thus indicating the intricate nature of LldD2’s evolutionary trajectory.

**Figure 1.**
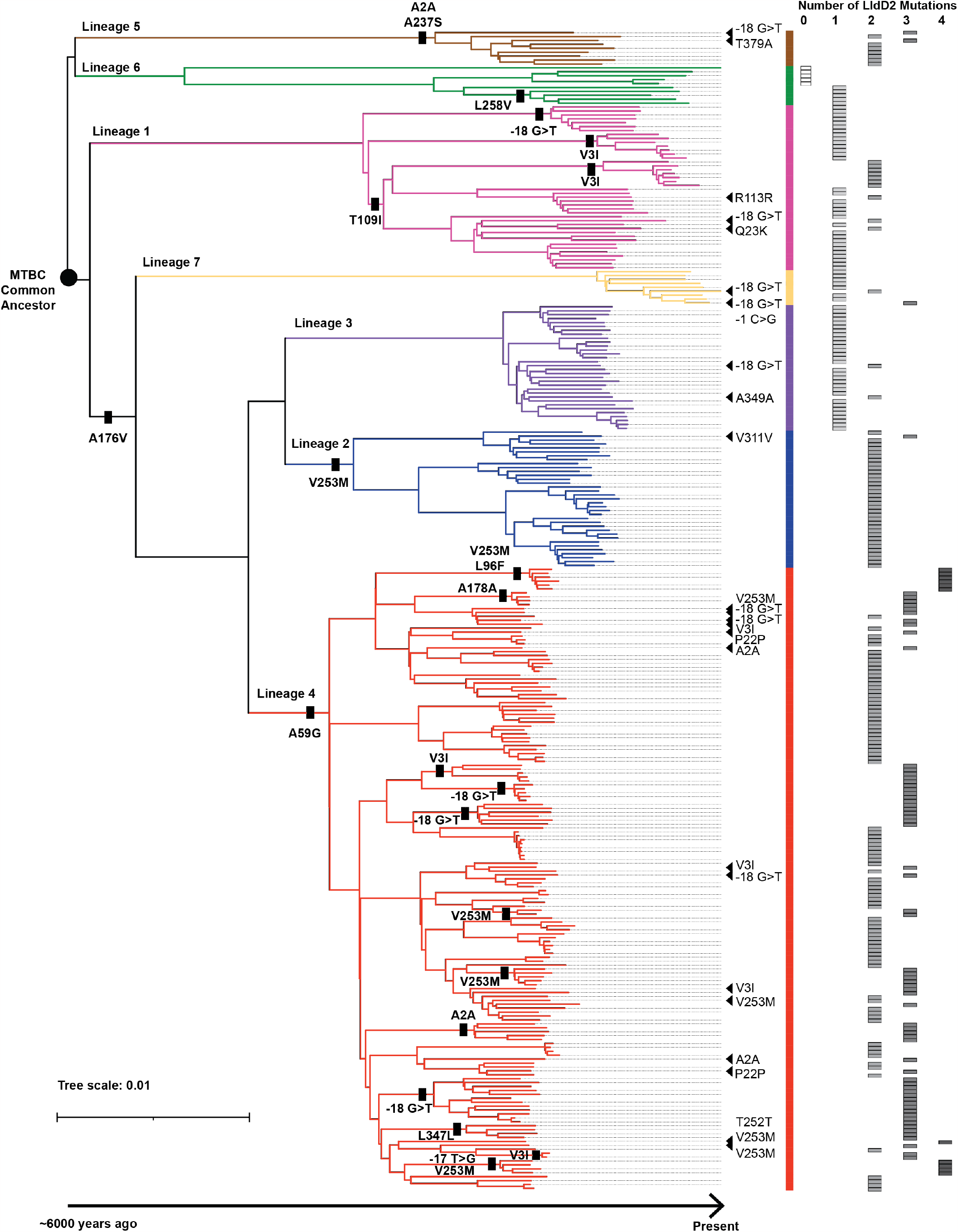
Stepwise evolution of LldD2. Phylogenetic tree of 296 *M. tuberculosis* and *M. africanum* clinical isolates representing the global diversity of the *Mycobacterium tuberculosis* complex (MTBC). The tree scale denotes number of mutations per site. Ancestrally derived lineage and sub-lineage defining *lldD2* mutations are indicated by a bar. Triangles denote *de novo* or homoplastic mutations. Total number of mutations per strain are tallied to the right of the phylogenetic tree.

**Figure 2.**
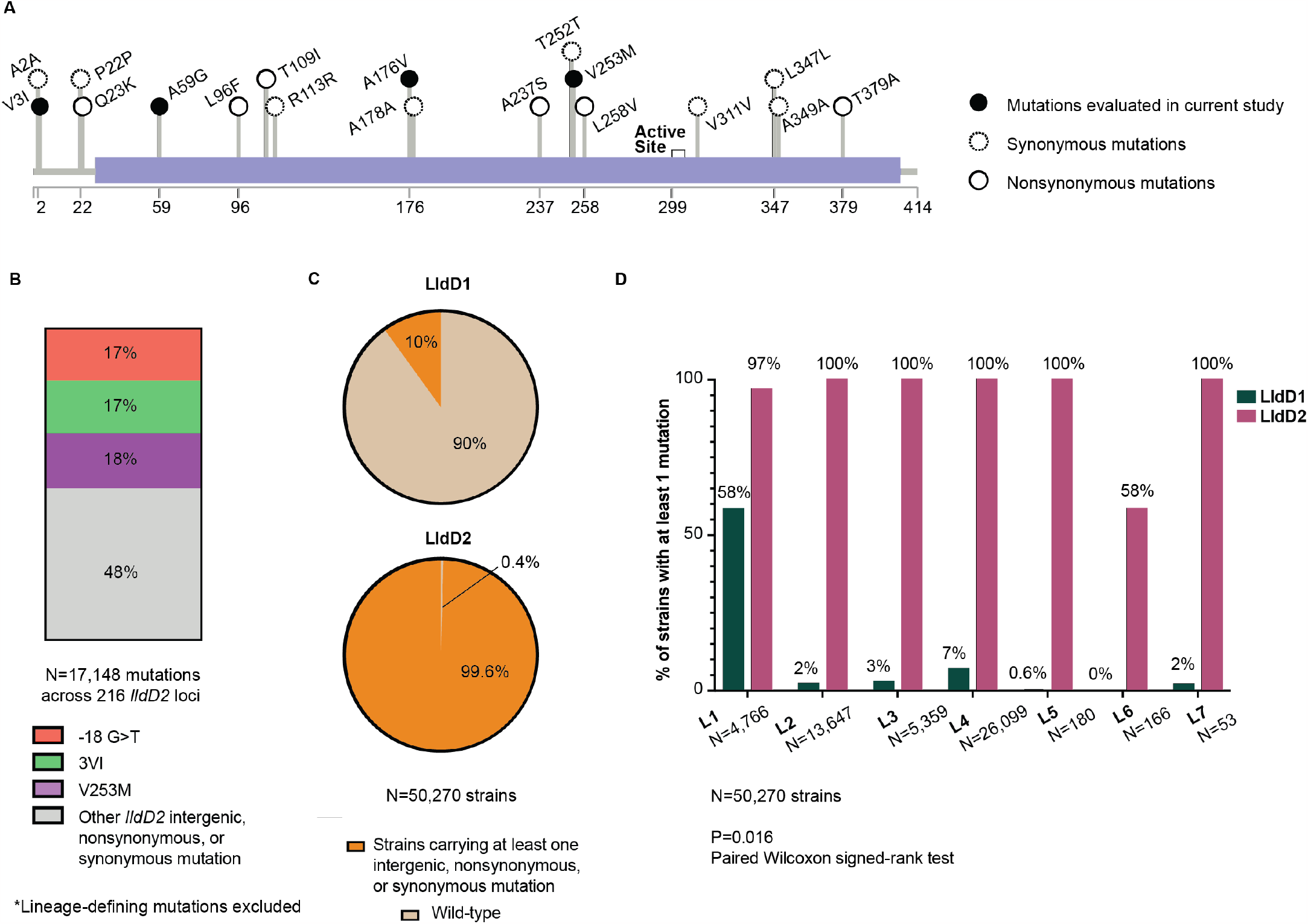
Prevalence of various LldD2 mutations. A. Schematic of LldD2 depicting the amino acid positions of the mutations listed in Fig. 1. The purple rectangle illustrates the FMN-dependent alpha-hydroxy acid dehydrogenase motif. Promoter mutations are not shown. B. Percentage of *lldD2* homoplastic or clade-defining mutations belonging to the indicated genotype. 17,148 mutations spanning 216 unique loci were identified across 50,270 strains. C. Percentage of strains carrying at least one of the indicated mutations in either LldD1 or LldD2. D. Comparison of the percentage of strains carrying at least one LldD1 or LldD2 mutation for each lineage.

The stepwise evolution of *lldD2* is especially striking in comparison to *lldD1*, which does not have LDH activity but is thought to participate in the oxidation of other small α-hydroxy acids (20). *LldD1* mutations have accumulated in only 10% of the 50,000 MTBC strains, and most of these variants are concentrated in L1 (Fig. 2C-D). However, 99.6% of MTBC strains carry at least one *lldD2* mutation (Fig. 2C). This distinct pattern of selection strongly suggests that natural selection has specifically targeted lactate metabolism.

To define the functional effects of the *lldD2* variants, we first identified clinical strains of Mtb with various *lldD2* mutations from a panel of strains isolated from TB patients in Ho Chi Minh City, Vietnam (30) (S1 Table). The clinical isolates belong to L1, L2, and L4, which are globally the three most prevalent lineages (31). We focused on the -18 G>T promoter mutation, V3I, and V253M because we identified both lineage/clade-defining and homoplastic occurrences of these variants in clinical strains (Fig. 1). We assessed Mtb growth in standard media (7H9), which contains glycerol as a carbon source, and defined media with lactate as the sole carbon source (7H12 – 0.2% L-lactate). After normalizing to growth in 7H9, we find that clinical isolates that carry the evolved *lldD2* alleles have increased growth in lactate media as compared to nearest-neighbor strains that only carry ancestral *lldD2* alleles (unpaired t-test, P=0.037) (Fig. 3A-B).

**Figure 3.**
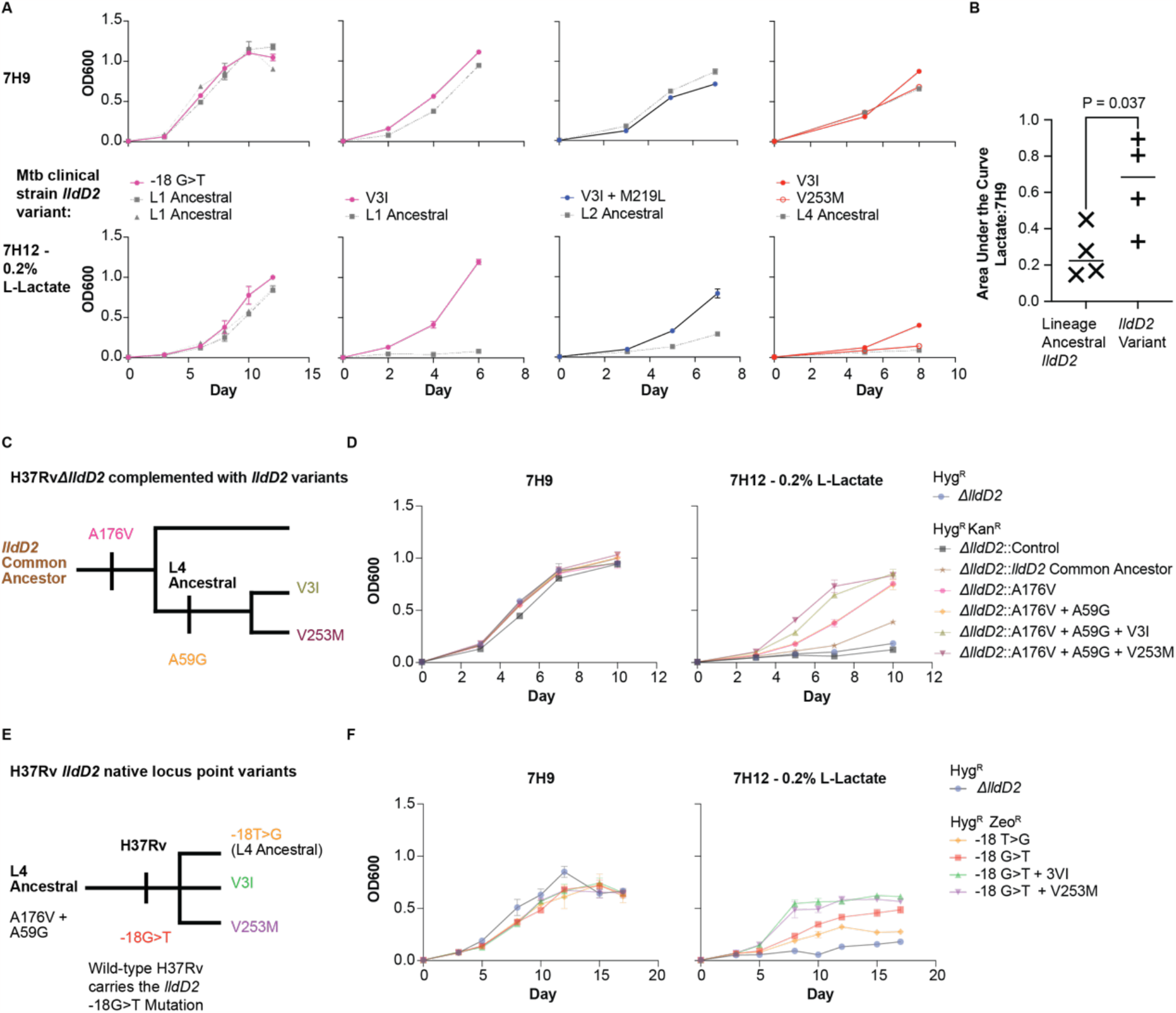
Ancestral and homoplastic mutations in *lldD2* augment Mtb fitness in lactate. A. Growth curves of Mtb clinical strains from L1, L2, and L4. Strains are grouped for comparison of the *lldD2* homoplastic mutation strains to closely related lineage ancestral strains. All cultures started at OD_600_ 0.005 at day 0. Triplicate replicates shown, error bars represent the standard deviation. B. Quantification of the area under the curves (AUCs) from A. The AUC in 7H12 – 0.2% L-lactate is normalized to that of 7H9 for a given strain. The AUC is averaged for the V3I and V253M L4 strains and to the two L1 ancestral strains that were paired to -18 G>T. P-value indicates results of an unpaired t-test. C and E. Schematic depicting the construction of recombinant *lldD2* strains. D and F. Growth curves of the strains indicated in C and E. Hyg^R^, Kan^R^, and Zeo^R^ refers to hygromycin, kanamycin, and zeocin resistance respectively. All cultures started at OD_600_ 0.005 at day 0. Triplicate replicates shown, error bars represent the standard deviation. Representative of two independent experiments.

We next constructed a *lldD2* deletion mutant in H37Rv, the L4 derived lab adapted strain. We complemented the mutant with a panel of *lldD2* alleles that recapitulate the evolution of *lldD2* in L4 from the inferred common ancestor sequence to the homoplastic variants present in L4 clinical strains (Fig. 1, Fig. 3C). In the engineered strains, expression of *lldD2* is driven by a constitutive promotor so the -18 G>T mutation was not assessed. In multiple carbon source media (7H9), all of the strains demonstrate similar growth patterns (Fig. 3D, fig. S1A). In lactate media, *ΔlldD2* and *ΔlldD2* complemented with a vector control (*ΔlldD2*::Control) do not grow appreciably. However, the addition of each successive *lldD2* mutation measurably augments Mtb growth in lactate, except for A59G for which we did not find a fitness benefit (Fig. 3D, fig. S1A).

Finally, we constructed isogenic allelic variants by introducing the point mutations within the native locus of *lldD2*. We constructed these variants in H37Rv, which contains the homoplastic - 18 G>T promoter mutation (Fig. 3E). Therefore, to create a strain expressing the L4 ancestral *lldD2* sequence, we introduced a -18 T>G mutation (Fig. 3E). We also introduced the V3I and V253M mutations into wild-type H37Rv (Fig. 3E). In 7H9, there were minimal quantitative differences in growth between strains (Fig. 3F, fig. S1B). However, in lactate media, we again see the strains expressing the evolved mutations grow faster than the strain expressing the L4 ancestral allele (−18 T>G) (Fig. 3F, fig. S1B).

### Homoplastic *lldD2* mutations alter gene and enzyme activity

To determine how the homoplastic *lldD2* mutations result in gain-of-function activity, we first assessed the effect of the promoter and early codon mutations on *lldD2* expression. We found that -18 G>T promoter mutation increases *lldD2* expression relative to the ancestral allele (ordinary one-way ANOVA, Dunnett’s multiple comparison test, P=0.011) (Fig. 4A). The addition of the V3I mutation (ordinary one-way ANOVA, Dunnett’s multiple comparison test, P=0.025) but not the V253M mutation (ordinary one-way ANOVA, Dunnett’s multiple comparison test, P=0.74) to the -18 G>T background also results in increased *lldD2* expression. To determine whether this results in increased LldD2 protein production, we constructed *M. smegmatis* (Msm) strains that express the first 60 amino acids of LldD2 fused to Renilla luciferase, with expression driven by the *lldD2* native promoter. Therefore, we could quantify luminescence as a proxy for LldD2 production. In this system, the -18 G>T mutation alone results in a minimal increase in LldD2 production compared to the L4 ancestral promoter (ordinary one-way ANOVA, Dunnett’s multiple comparison test, P=0.48), but consistent with the gene expression data, both the V3I and the -18 G>T + V3I variants significantly increase protein production (ordinary one-way ANOVA, Dunnett’s multiple comparison test, P=0.0002 and P<0.0001 respectively) (Fig. 4B).

**Figure 4.**
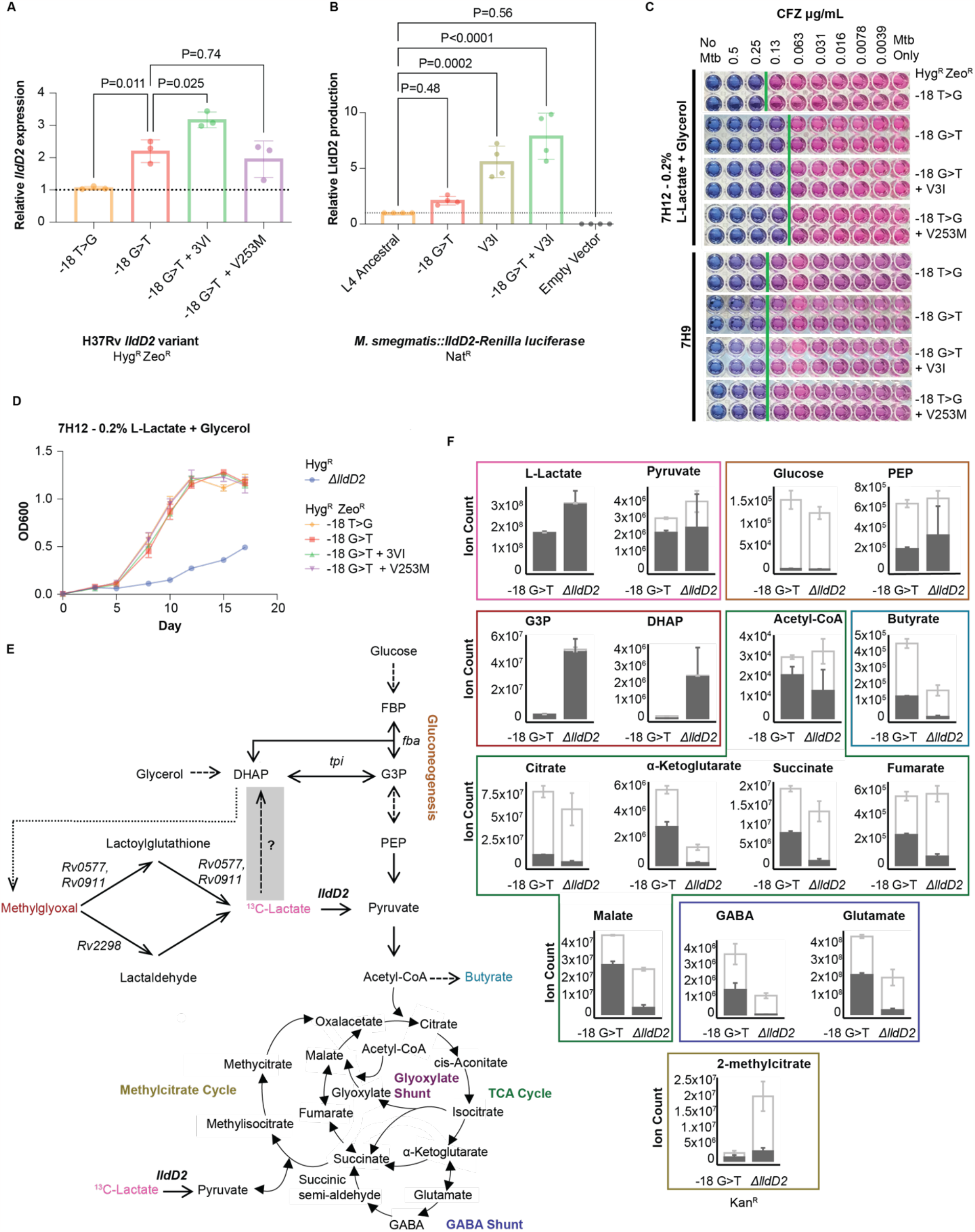
*lldD2* homoplastic mutations modulate gene and enzyme activity. A. *lldD2* expression as measured by qPCR. Gene expression is normalized to that of -18 T>G. Each dot represents average of technical replicates from one of three independent experiments. Error bars indicated standard deviation. P-value indicates results of an ordinary one-way ANOVA test with Dunnett’s multiple comparison correction. B. LldD2 production as measured by a Renilla luciferase assay. Luminescence normalized to that of the Msm strain expressing the L4 ancestral version of *lldD2*. Each dot represents average of technical replicates from an independent experiment, error bars represent the standard deviation. P-value indicates results of an ordinary one-way ANOVA test with Dunnett’s multiple comparison correction. Nat^R^ refers to nourseothricin resistance. C. Alamar blue MIC assay of clofazimine (CFZ). Green bars indicate the MIC. Representative of three independent experiments. D. Growth curves of the indicated strains. All cultures started at OD_600_ 0.005 at day 0. Triplicate replicates shown, error bars represent the standard deviation. Representative of two independent experiments. E. Schematic of ^13^C-Lactate metabolic flux assay. Grey box indicates a proposed pathway. Dotted arrow represents the spontaneous conversion of DHAP to methylglyoxal. Dashed arrows represent abbreviated steps. F. ^13^C-Lactate metabolic flux assay results. Error bars represent the standard deviation of three technical replicates. Purple indicates labeled carbon ions, white represents total carbon ions.

LldD2 mediates the oxidation of lactate into pyruvate, which is a quinone-dependent reaction (20). Given that we have thus far demonstrated that the homoplastic *lldD2* mutations are gain-of-function, we hypothesized that the variants might affect the minimum inhibitory concentration (MIC) of clofazimine (CFZ). CFZ binds to NDH-2, the main NADH:quinone reductase of the Mtb electron transport chain, with higher binding affinity than menaquinone; oxidation of CFZ results in reactive oxygen species (ROS) production which ultimately leads to Mtb killing (32). Accordingly, CFZ sensitivity can serve as a probe of quinone dependent enzyme activity. In 7H9, all strains have the same MIC to CFZ (Fig. 4C). To test the effect of CFZ in the presence of lactate, we used 7H12 containing 0.2% glycerol and L-lactate. We included glycerol so that the *lldD2* variants would grow at the same rate even with lactate present (Fig. 4D, fig. S1C). Mtb strains expressing homoplastic *lldD2* variants demonstrate a 2-fold lower MIC compared to the - 18 T>G strain expressing the L4 ancestral allele (Fig. 4C). All strains demonstrated the same susceptibility to trifluoperazine, which targets NDH-2 via a different mechanism of action (33) (fig. S2). These data suggest that *lldD2* gain-of-function variants alter quinone cycling during lactate metabolism.

In these studies, the *ΔlldD2* strain demonstrated markedly slower growth in the media condition containing both lactate and glycerol (Fig. 4D, fig. S1C). Billig *et al*. showed that *lldD2* is necessary for growth with lactate as the sole carbon substrate (20), but our data imply that growth in lactate is deleterious to Mtb without a functional lactate dehydrogenase, even in the presence of another carbon source. Indeed, *ΔlldD2* does not exhibit a growth defect when glycerol is the sole carbon source (fig. S3). Thus, we postulated that LldD2 has a detoxication function in addition to its role in anaplerosis. Consistent with this, Pethe *et al*. suggested that LldD2 participates in the detoxification of methylglyoxal, a toxic byproduct that is formed spontaneously as a result of the metabolism of triose phosphates such as dihydroxyacetone phosphate (DHAP) and glycerol-3-phosphate (G3P) (Fig. 4E) (34). We reasoned that impaired growth of the *ΔlldD2* strain may be due to the disruption of methylglyoxal detoxification (Fig. 4E).

To test this model, we constructed a pair of *ΔlldD2* and -18 G>T strains in the same parental genetic backgrounds (fig. S4) (S1 Table). We performed metabolic flux assay after 8 hours of growth on ^13^C-lactate and glycerol and observed markedly more ^13^C-labeled DHAP and G3P in the *ΔlldD2* strain compared to the -18 G>T strain (Fig. 4F) (S2 Table). DHAP and G3P were nearly completely labeled in the *ΔlldD2* strain, indicating that these metabolites were the products of ^13^C-lactate metabolism. We also observed significant ^13^C-incorporation of phosphoenolpyruvate (PEP) but not glucose, which is indicative of glycolytic but not gluconeogenic flux of lactate under this growth condition (Fig. 4E-F). We found labeled lactate and pyruvate in both -18 G>T and Δ*lldd2* strains (Fig. 4F). Together, these data suggest that without LldD2 activity, ^13^C-lactate buildup favors DHAP and G3P formation. This drives glycolytic flow to pyruvate, but also mediates methylglyoxal formation, which leads to the growth impairment of the *ΔlldD2* strain in the presence of lactate (Fig. 4E-F). This implies a metabolic pathway exists in which lactate is a precursor of DHAP or G3P synthesis.

We completed the same experiment with the Mtb strains expressing the evolved *lldD2* point mutants. We compared the ancestral -18 T>G strain to -18 G>T, -18 G>T + V3I, and -18 G>T + V253M variants and similarly found increased G3P labeling in the ancestral *lldD2* strain compared to those carrying evolved variants (fig. S5). Together, the growth curve and metabolic flux data provide experimental evidence for the proposed role of LldD2 in both incorporating lactate into central carbon metabolism and ameliorating lactate toxicity by reducing methylglyoxal accumulation. Therefore, mitigating methylglyoxal toxicity may be a contributing selective force driving gain-of-function *lldD2* diversification.

### Lactate rewires transcriptomic networks involving Mtb cell wall lipids and virulence systems

We next sought to identify downstream effects of *lldD2* variation during Mtb metabolism of lactate. We performed genome wide gene expression analysis of the -18 G>T and -18 T>G strains after 6 hours in 7H9 or in 7H12 with lactate as the sole carbon source. A principal component analysis (PCA) of the technical replicates based on reads demonstrated that lactate drives significant differences in Mtb gene expression (Fig. 5A). In 7H9, just seven genes are significantly upregulated (log_2_ fold change (L2FC) > 0.6 and P < 0.01) in the -18 G>T strain compared to the -18 T>G strain (Fig. 5B, S3 Table). Of these, only three gene have L2FC >1: *lldD2*, Rv1871c, and Rv1870c, consistent with our directed analysis (Fig. 3A) and operonic expression of Rv1872c-Rv1870c (Fig. 5B, S3 Table).

**Figure 5.**
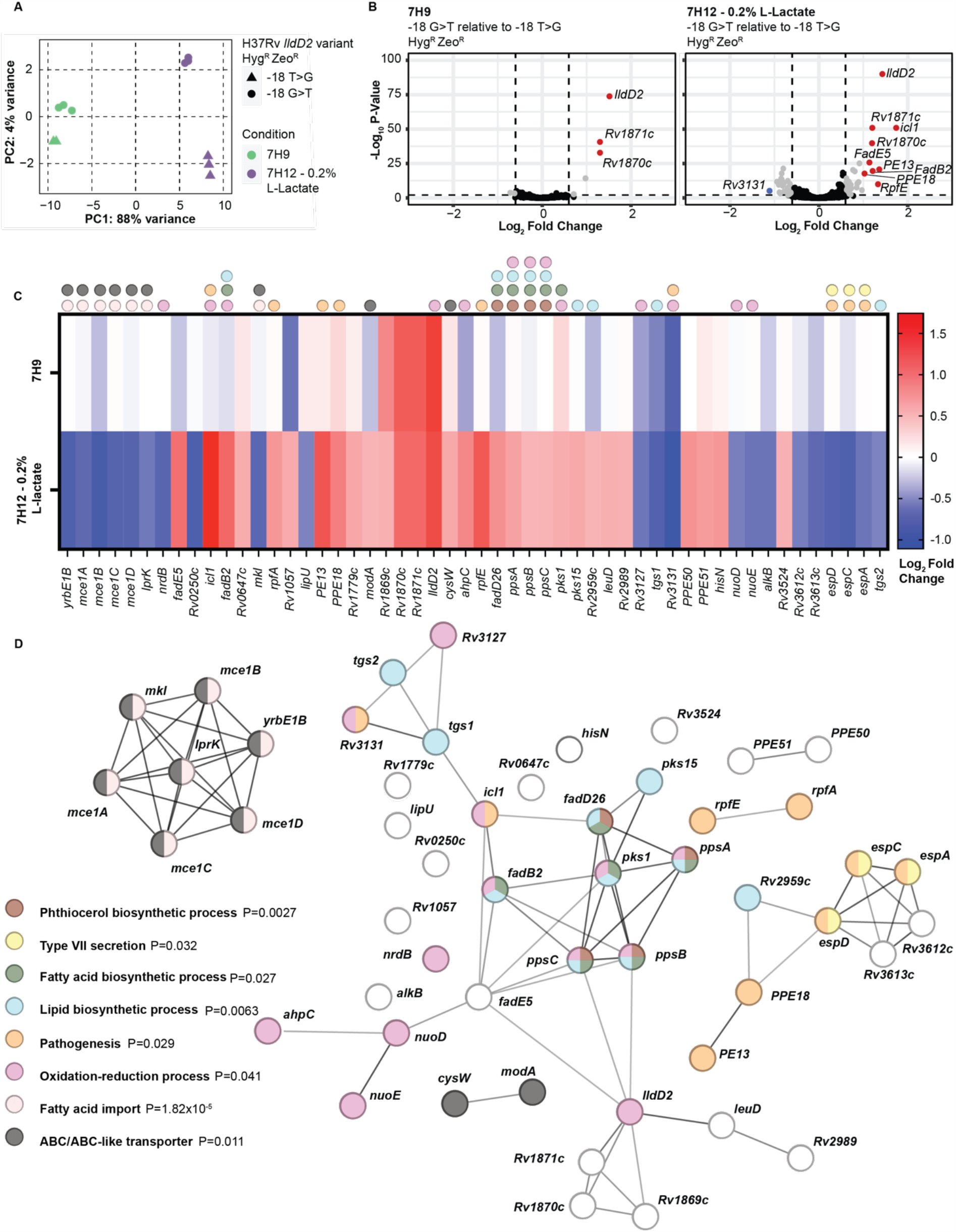
Lactate and *lldD2* variants remodel the Mtb virulence transcriptome. A. Principal component analysis of the RNA-seq technical replicates based on normalized counts. B. Volcano plot of the differential expression (DE) analysis. Grey indicates DE genes with log_2_ fold change (L2FC) greater than 0.6 or less than -0.6 (P<0.01), red indicates DE genes with a L2FC greater than 1, blue indicates DE genes with a L2FC less than -1. There are 7 DE genes in 7H9 and 52 DE genes in lactate. C. Heatmap of the 52 DE genes from the lactate condition, 7H9 shown for comparison. The genes are ordered according to chromosome position. Dots denote genes involved in the pathways shown in D. D. STRING protein-protein interaction network of the 52 DE genes from the lactate condition. P-value indicates the significance of the Gene Ontology, KEGG, or STRING pathway enrichment after multiple test correction.

In the lactate growth condition, 52 genes were differentially expressed (L2FC greater than 0.6 or less than -0.6, P<0.01) comparing ancestral -18 G>T to -18 T>G (Fig. 5B-C) (S3 Table). We utilized this gene set to create protein-protein interaction network based on the STRING database and performed functional enrichment analysis according to Gene Ontology biological processes (36), STRING network clusters, and KEGG pathways (37). This gene set has significantly more interactions compared to a sample randomly selected from the genome (P=2.21x10^−11^), which is indicative of a coordinated biological response (35). During lactate metabolism, the -18 G>T mutation modulates expression of gene programs largely involved in fatty acid metabolism and virulence. This gene set is enriched for genes that coordinate fatty acid import (P=1.82x10^−5^), notably the *mce1* operon, which is downregulated in the strain with the evolved allele (Fig. 5C-D). Differently expressed genes are also enriched for mediators of fatty acid (P=0.027), lipid (P=0.0063), and phthiocerol biosynthetic processes (P=0.0027) such as *fadB2, fadD26, pks1, pks15*, and *ppsA-C*, which are all upregulated (Fig. 5C-D) (38-41). We also find upregulation of *icl* which encodes an isocitrate lyase (Fig. 5C-D); Serafini *et al*. found similar transcriptional changes in Mtb grown on lactate and also showed that *icl* is essential for Mtb growth in lactate (42). In addition to *icl*, this gene set is enriched with other genes implicated in oxidation-reduction processes like electron transport components *nuoD* and *nuoE*, which are both downregulated in the strain carrying the evolved allele (Fig. 5C-D) (43).

Type VII secretion components (P=0.032) are also enriched in this gene set. We find that the *espACD* operon is downregulated in Mtb with the evolved -18 G>T allele compared to -18 T>G ancestral allele in lactate containing media (Fig. 5C-D). This operon is essential for secretion of the virulence effectors EsxA and EsxB via the ESX-1 Type VII secretion system (44, 45).

Serafini *et al*. also observed downregulation of *espACD* expression in H37Rv (which expresses the derived *lldD2* -18 G>T allele) grown on lactate compared to glucose (42). Other genes involved in pathogenesis (P=0.029) are upregulated including those encoding PE13, T-cell receptors specific to which are associated with TB disease control (46), and PPE18, which is a component of the M72/AS01_E_ TB vaccine candidate (47). These data indicate that lactate metabolism is not just feeding anaplerotic needs but is a host-relevant signal that the bacterium responds to by changing expression of key virulence systems.

## Discussion

Previous work determined that *lldD2* is undergoing host-derived positive selection and identified lineage and sub-lineage defining mutations in *lldD2* (8,10,21). We expanded this analysis to demonstrate the stepwise addition of *lldD2* mutations that occurred over evolutionary history and showed this augments Mtb replicative fitness during lactate growth. A very simple model for the selective advantage conferred by *lldD2* evolution is that that lactate is an abundant carbon source in some host environments, and Mtb is simply tuning lactate fitness via the evolution of *lldD2* to increase anabolic capacity.

However, our data support and expand a more complex model proposed by Pethe *et al*. in which a detoxification pathway exists between lactate and methylglyoxal, a byproduct produced spontaneously during glycolysis and lipid metabolism (34). Consistent with this model, we find that DHAP and G3P, precursors to methylglyoxal, accumulate in the *ΔlldD2* strain when grown in the presence of lactate and glycerol. *LldD2* gain-of-function variants mitigate this accumulation. Interestingly, LDH knockout or deficiency has also been associated with an increase in G3P in *Drosophila melanogaster* larvae, cancer cell lines, and in human patients (48-51). This suggest to us that detoxification may also be a significant selective pressure in addition to the anaplerotic importance of LldD2.

Finally, transcriptomic analysis demonstrates that *lldD2* mutations have effects beyond lactate anaplerosis. Evolutionary tuning of lactate metabolism affects genes involved in phthiocerol synthesis – implicated in macrophage evasion and immunopathology (52) – as well as ESX1 function. This suggests that Mtb modifies its virulence strategies when exposed to lactate and that the evolution of *lldD2* could be driven in part by cell wall and/or virulence remodeling.

Lactate production is induced by infected macrophages after stimulation with IFNγ (25), which suggests lactate is an immune-regulated carbon substrate. Mtb may utilize lactate as a signal of the host immune state and adjust growth and virulence accordingly. This would suggest that lactate is an ancient selective pressure that is still relevant today as *lldD2* diversification is currently ongoing (8). Assuming that lactate has always been a relevant feature of the host environment, it is unclear why optimized *lldD2* function did not evolve long ago. This could simply reflect the stochasticity of evolution. The Mtb strain carrying the most ancestral allele of *lldD2* is significantly impaired during growth in lactate, which may suggest that LldD2 once mediated enzyme activity distinct from lactate oxidation. Over time, stochastic acquisition of mutations that enabled lactate oxidation may have tuned the ability of LldD2 to function as a LDH, and these mutations were selected for their pleiotropic effects on Mtb fitness during lactate growth. There is precedence for the hypothesis that LldD2 was derived from an enzyme with a different function; a previous study demonstrated that SNPs can confer LDH activity to a bacterial glycerol dehydrogenase (53). Alternatively, the availability of human lactate during infection could be dynamic, which would force Mtb to adapt. This change in host environment could be shaped by shifts in immune state, perhaps from BCG vaccination, altered nutritional state or altered transmission pressures – modulating either lactate exposure or the optimal response to this host cue.

In all, we provide experimental evidence that ancient lineage and sub-lineage defining mutations and more contemporary homoplastic mutations in *lldD2* reflect stepwise gain-of-function. This work has provided novel insight into Mtb adaption to host stress, and as a result, elucidated the intersection of metabolic remodeling and virulence.

---

**Supplementary Figure 1.**
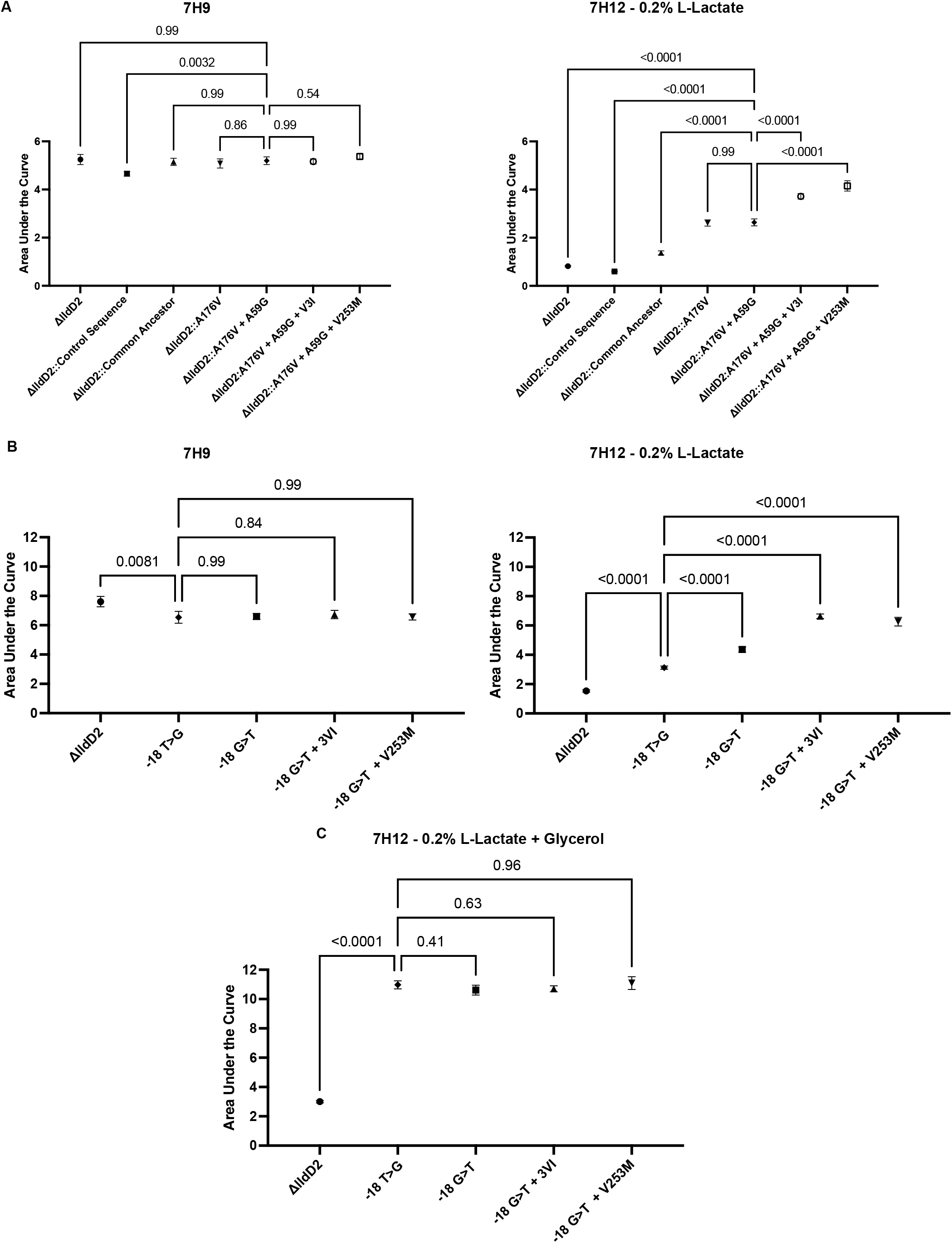
Comparison of the area under the curve for the growths curves shown in Fig. 3D (A), Fig. 3F (B), and Fig. 4D (C). Three replicates are shown, error bars indicate the standard deviation. P values indicate the results of an ordinary one-way ANOVA with Dunnett’s multiple comparison test. Representative of two independent experiments.

**Supplementary Figure 2.**
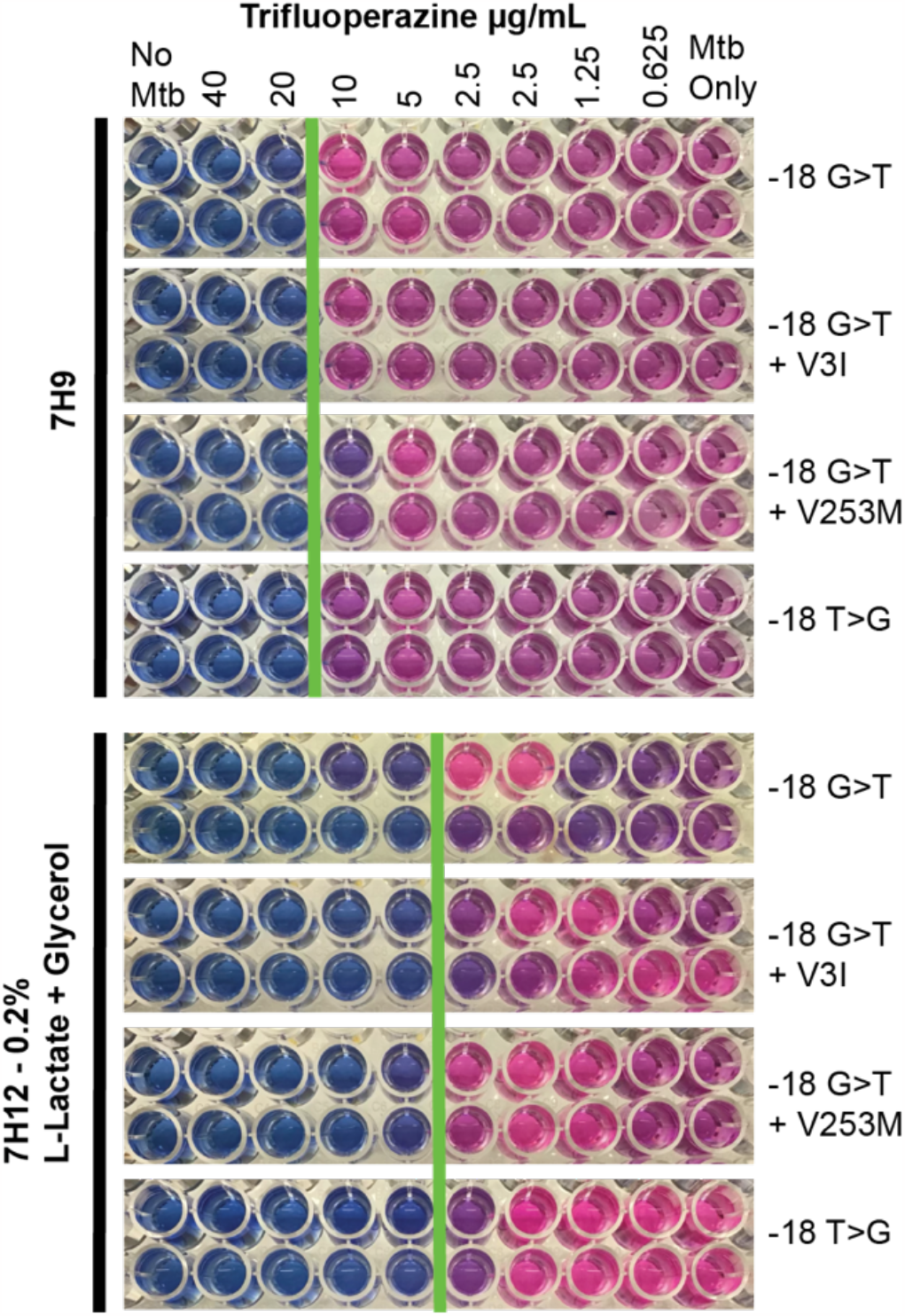
Minimum inhibitory concentration (MIC) assay results. Technical replicates for each strain are shown, the data is representative of two independent experiments. Green bars indicate the MIC.

**Supplementary Figure 3.**
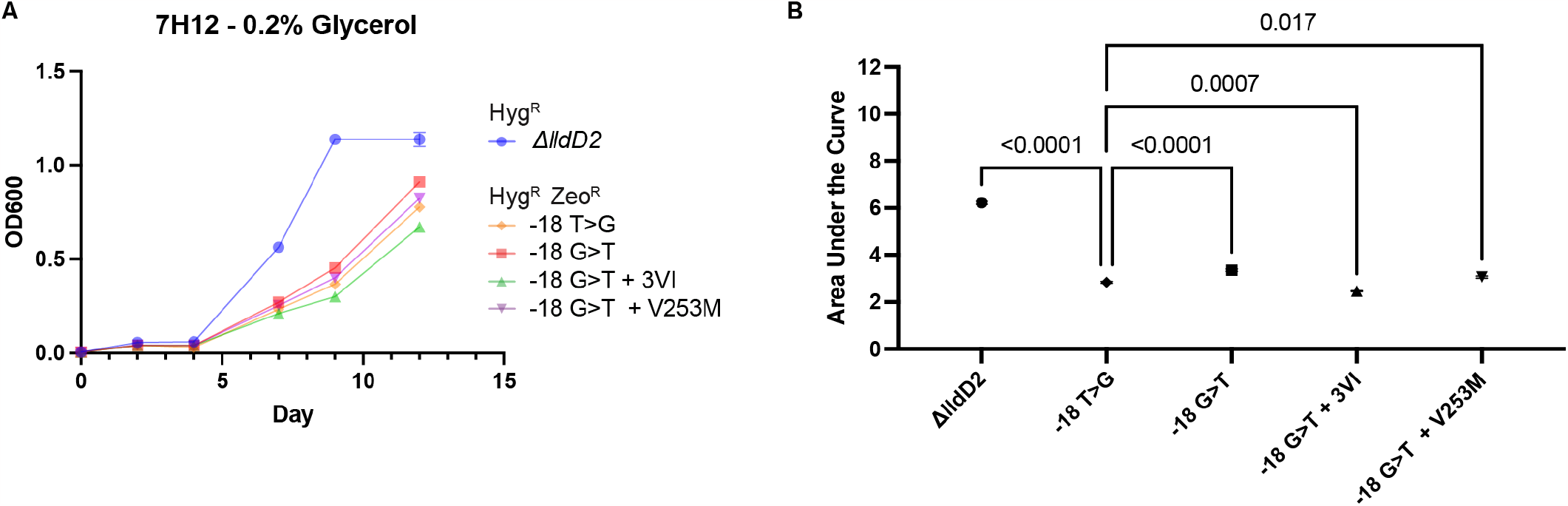
A. Growth curves of the *lldD2* allelic variants with glycerol as the sole carbon source. All cultures started at OD_600_ 0.005 at day 0. Triplicate replicates shown, error bars represent the standard deviation. Representative of two independent experiments. B. Area under the curve analysis of the growth curves shown in A. Three replicates are shown, error bars indicate the standard deviation. P-values indicate the results of an ordinary one-way ANOVA with Dunnett’s multiple comparison test.

**Supplementary Figure 4.**
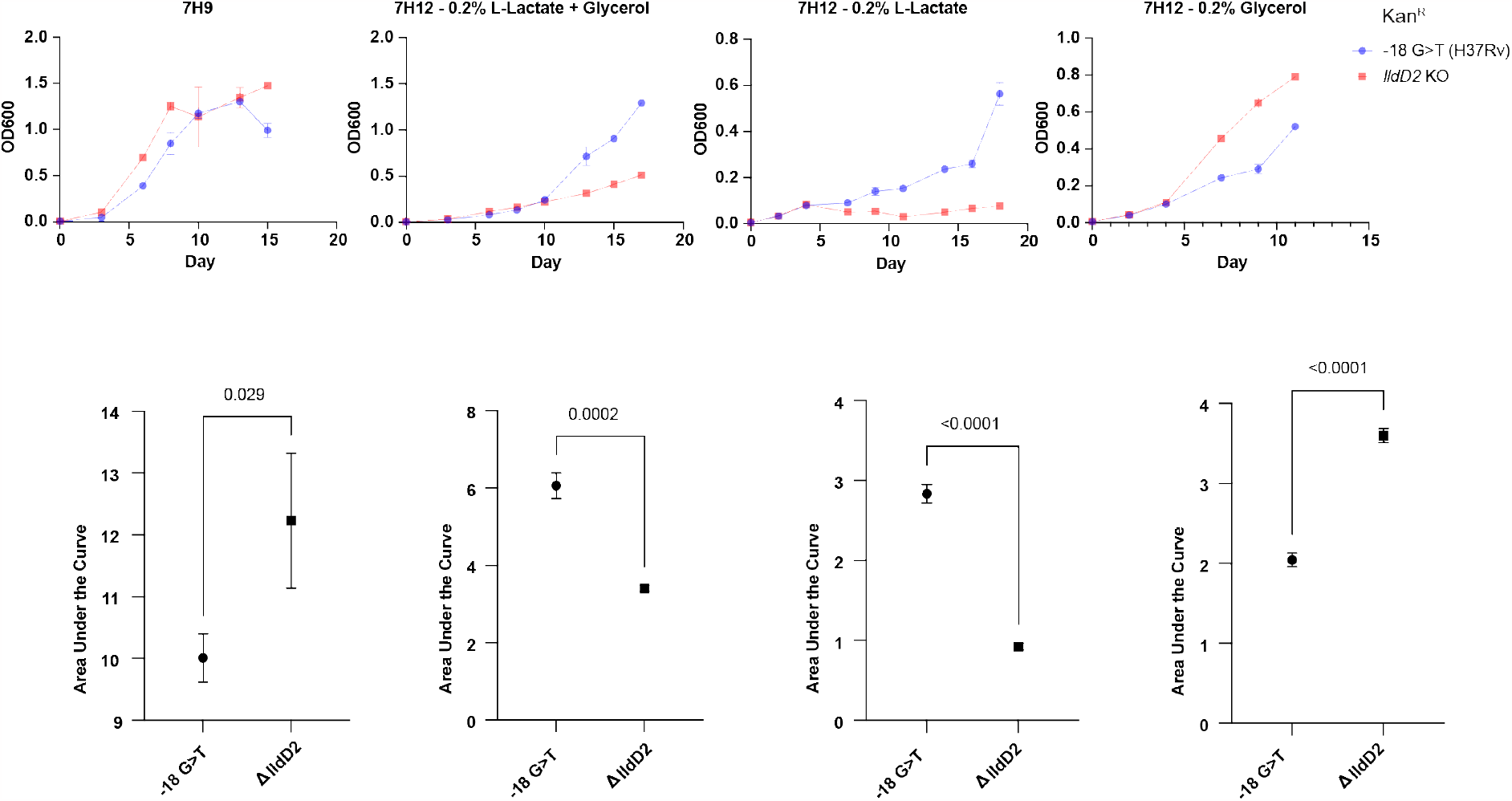
Growth curves and corresponding area under the curve analysis for the strains utilized for the metabolic flux assay shown in Fig. 4E. All cultures started at OD_600_ 0.005 at day 0. Triplicate replicates shown, error bars represent the standard deviation. H37Rv carries the -18 G>T *lldD2* mutation. Kan^R^ refers to kanamycin resistance. P-values indicate the results of unpaired t-tests. Representative of two independent experiments.

**Supplementary Figure 5.**
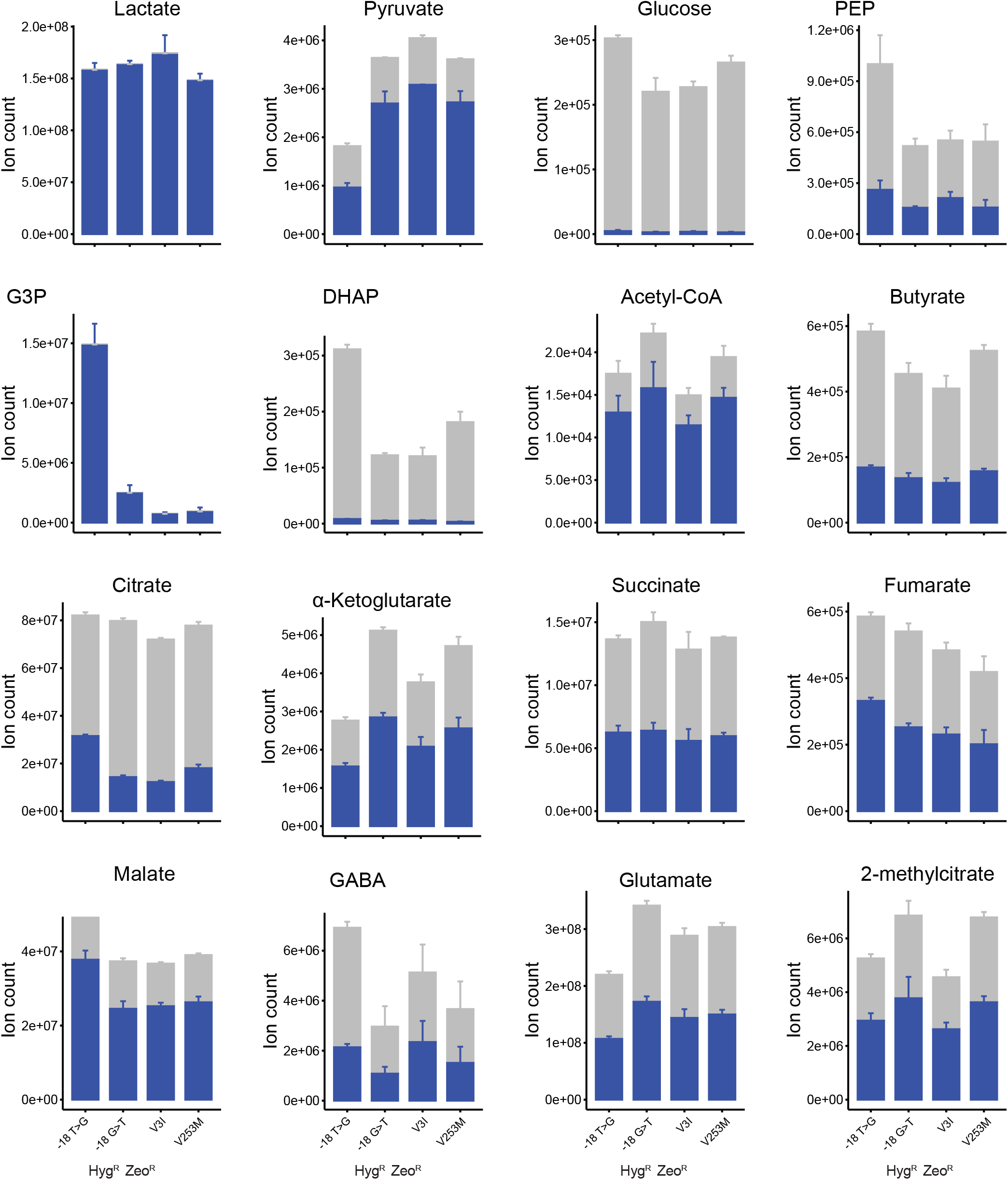
^13^C-lactate metabolic flux analysis results. The error bars represent the standard deviation of three technical replicates. Blue indicates labeled carbon ions, grey represents total carbon ions.

