## Supplementary material for "Ongoing evolution of the *Mycobacterium tuberculosis* lactate dehydrogenase reveals the pleiotropic effects of bacterial adaption to host pressure": Methods

***lldD2* mutant strain construction**

We completed oligo-mediated recombineering according to the protocol as described (1,2) in the Mtb H37Rv reference strain to introduce the V3I and V263M mutations into the native locus of the coding region of *lldD2* and to revert the -18G>T mutation acquired by H37Rv to the ancestral allele. We retained strains that failed to incur *lldD2* mutations for use as H37Rv controls.

*ΔlldD2* was constructed in H37Rv according to the methodology detailed by Griffin *et al* (3). We then complemented *ΔlldD2* with a series of different *lldD2* sequences that recapitulate the evolutionary history of the gene. We also complemented *ΔlldD2* with the first 300 base pairs upstream of *lldD2*, this null sequence serves as a negative control. Complementation was accomplished by transformation with a plasmid carrying a kanamycin resistance cassette and a *lldD2* sequence of interest that integrates at the L5 phage site. *LldD2* was expressed under the direction of the pUV15-TetON promoter (4); because the plasmid we used lacks the repressor TetR, *lldD2* is constitutively expressed.

For Fig. 4E, we also cured the hygromycin cassette from the *lldD2* locus of the *ΔlldD2* strain (3). The cassette was removed by transforming *ΔlldD2* with a plasmid constitutively expressing a Cre recombinase that executes excision between the loxP sites. As an empty vector control, the H37Rv strain used to construct *ΔlldD2* was transformed with the same plasmid. We completed Sanger sequencing of the transformants to confirm loss of the hygromycin cassette and tested for susceptibly to hygromycin by inoculating the strains in 10 mL of 7H9 media (7H9 salts with 0.2% glycerol, 10% OADC supplement, 0.05% Tween80) and 50 µg/mL of hygromycin and using a spectrophotometer to monitor growth via OD_600_ for seven days.

The Msm strains expressing *lldD2* were constructed in the reference strain mc^2^155. We transformed mc^2^155 with a Giles integrating plasmid carrying a nourseothricin resistance cassette, a sequencing corresponding to 200 base pairs upstream of *lldD2*, and the *lldD2* sequences of interest encoding the first 60 amino acids of LldD2 with the C-terminus fused to Renilla luciferase. With the inclusion of the upstream region, *lldD2* expression is driven by the native promoter. The plasmid is modified from Rock *et al* (5). See S1 Table for a list of all strains used in the study.

**Mtb clinical strains**

The clinical strains utilized in this study are part of a larger collection of 1,635 samples originally isolated from patient sputum in Ho Chi Minh City, Vietnam and published by Holt *et al* (6). Our group transformed a subset of these strains with a L5-intergrating plasmid carrying a kanamycin resistance cassette and a unique genetic barcode for a separate analysis (7). These barcoded clinical strains were used to conduct the growth curves in the current analysis. S1 Table lists the *lldD2* genotype and accession numbers for these strains.

**Phylogenetic tree and *lldD2* mutation annotation**

The phylogenetic tree depicting the global diversity of MTBC strains was adapted from the figure originally published by Liu *et al* (8). The phylogenic tree was designed using iTOL (version 6.6) (9). The *lldD2* SNPs were called in reference to the inferred ancestral genome of the MTBC most recent common ancestor (10). These mutations were utilized to annotate the LldD2 diagram using the lollipops package (v1.6.0) (11). The *lldD1* and *lldD2* mutation burden figures in Fig. 2B-D were based on the sequences from the ~50,000 clinical isolates analyzed by Liu *et al* (2).

**Growth curves**

The growth curves with the recombinant Mtb strains and the clinical isolates were completed by diluting mid-log phase cultures (grown in 7H9 at 37° C with shaking) to an OD_600_ of 0.005 in 10 mL of the indicated media that included 50 µg/mL hygromycin, 20 µg/mL kanamycin, and/or 20 µg/mL zeocin. 7H12 consists of 7H9 salts, 0.1% casamino acids, and 0.05% tyloxapol. Carbon sources include 0.2% w/v lactate, 0.2% v/v glycerol, or 0.1% w/v lactate and 0.1% v/v glycerol (0.2% lactate + glycerol). Each strain was inoculated in three separate inkwells per media condition to serve as replicates. We measured the OD_600_ at the indicated time points after inoculation. We performed growth curves independently twice for each set of strains.

***lldD2* expression analyses**

We completed qPCR to measure *lldD2* expression in the point mutant strains. To do this, we inoculated each strain into 10 mL of 7H9 + 50 µg/mL hygromycin and allowed them to grow at 37° C with shaking to mid-log phase. In triplicate, we diluted each strain into 10 mL of 7H9 so they would reach an OD_600_ of 0.5 after 3 days of additional growth. Afterwards, we spun down the cultures, resuspended in 1 mL of TRIzol (Invitrogen, Waltham, MA, USA), and completed bead beating to lyse the cells. We added 30% total volume of chloroform then performed RNA extraction and cleanup using the RNA Clean & Concentrator kit (Zymo Research, Irvine, CA, USA). We obtained cDNA using random hexamers with SuperScript IV reverse transcriptase kit (Thermo Fisher Scientific, Waltham, MA, USA). We then performed qPCR to quantify transcript abundance. *LldD2* expression was normalized to *sigA* expression. We utilized the following primers: *sigA* 5ʹ-CAAGTTCTCCACCTACGCTAC-3ʹ and 5ʹ-GTTGATCACCTCGACCATGT-3ʹ; *lldD2* 5ʹ- CCGCGACATCGAGTTTCACCCG-3ʹ and 5ʹ- GGATGGACATCCCGTTGCGGACATC-3ʹ. We completed three independent rounds of culturing, RNA extraction, and qPCR.

**Renilla luciferase assay**

To quantify LldD2 production, we performed a Renilla luciferase assay using the Msm LldD2-Renilla luciferase fusion constructs (5). Each Msm strain was cultured to mid log phase in 5 mL of 7H9 + ADC (7H9 salts with 0.2% glycerol, 10% ADC supplement, 0.05% Tween80) plus 25 µg/mL nourseothricin at 37° C with shaking. In triplicate, the cultures were then back diluted into 7 mL of 7H9 + ADC and 25 µg/mL nourseothricin such that they would reach log phase growth after 24 hours of growth. We measured the OD_600_ of the cultures and harvest 4 OD_600_ units of cells by pelleting the samples. After removing the supernatant, we completed the assay according to protocol (Renilla Luciferase Assay System, Promega, Madison, WI, USA). We performed the assay in 96-well Costar white plates (Corning, Corning, NY, USA) and quantified luciferase fluorescence with the VarioSkan Flash plate reader (Thermo Fisher Scientific). The Renilla assay was performed independently four times.

**Minimum inhibitory concentration assay**

The minimum inhibitory concentration (MIC) for clofazimine (CFZ) and trifluoperazine was assessed with an alamar blue reduction assay, which we performed as described but without shaking and with use of the indicated media (12). The MIC was determined as the lowest concentration of antibiotic that inhibited the alamar blue reagent (BioRad, Hercules, CA, USA) from transitioning from blue to purple or pink after 3 days of incubation. We performed the CFZ MIC assay three independent times and the trifluoperazine assay two independent times.

**Lactate metabolic flux assay**

We cultured the point mutant strains and the cured *ΔlldD2* strains to mid-log phase in 7H9 with the appropriate antibiotics, then back diluted the strains so they would read an OD_600_ of ~1 two days later at 37° C with shaking. We collected the cells via vacuum filtration by applying 1 mL of culture to a 0.22 µm mixed cellulose filter membrane (MilliporeSigma, St. Louis, MO, USA) affixed to a filter membrane holder (MilliporeSigma). We collected three samples per strain. The Mtb-laden filters were placed on 7H10 solid media (7H10 salts, 0.1% casamino acids, 0.1% v/v glycerol, 0.1% w/v L-lactate, and the appropriate antibiotic) and incubated for 7 days at 37° C to allow for biomass formation. We then pulsed with ^13^C-labelled lactate by transferring the Mtb biomass filters onto 7H10 solid media with 0.1% v/v glycerol, 0.1% w/v ^13^C-laballed lactate, and the appropriate antibiotic for 8 hours. Afterwards we quenched metabolism by placing each filter into a 40:40:20 solution of acetonitrile:methanol:water chilled with dry ice for 1 minute. The transfer buffer, Mtb biomass, and filter were then transferred into bead-beating tubes filled with 0.1 mm Zirconia/silica beads for metabolite extraction. We completed bead-beating 6 times for 30 seconds at 6 m/s with intermittent cooling down for 1 minute on ice. After centrifugation, we double-filtered the samples with Spin-X centrifuge 0.22 µm cellulose-acetate filter tubes (Corning) for removal out of the BSL3 laboratory. We used the Pierce BCA protein assay kit (Thermo Fisher Scientific) to measure residual protein in the samples in order to normalize by cell abundance in downstream analyses. Sample were stored at -80° C.

Metabolite extract samples were analyzed using a quadrupole-orbitrap mass spectrometer coupled with hydrophilic interaction chromatography (HILIC) as the chromatographic technique. Chromatographic separation was achieved on an XBridge BEH Amide XP Column (2.5 µm, 2.1 mm × 150 mm) with a guard column (2.5 µm, 2.1 mm X 5 mm) (Waters, Milford, MA, USA). For the gradient, mobile phase A was water:acetonitrile 95:5, and mobile phase B was water:acetonitrile 20:80, both phases containing 10 mM ammonium acetate and 10 mM ammonium hydroxide. The linear elution gradient was: 0 ~ 3 min, 100% B; 3.2 ~ 6.2 min, 90% B; 6.5. ~ 10.5 min, 80% B; 10.7 ~ 13.5 min, 70% B; 13.7 ~ 16 min, 45% B; and 16.5 ~ 22 min, 100% B, with a flow rate of 0.3 mL/ min. 5 µL of samples were injected using the autosampler at 4˚C. Needle wash was applied between samples using methanol:acetonitrile: water at 40:40:20. The mass spectrometer used was Q Exactive HF (Thermo Fisher Scientific), and scanned from 70 to 1000 *m/z* with switching polarity. The resolution was 120,000. Metabolites were identified based on accurate mass and retention time using the EI-Maven (Elucidata, Cambridge, MA, USA) with an in-house library. Correction of isotope-labeled lactate or ^13^C-natural abundance was performed in R using package AccuCor (13). We calculated the relative abundance of isotopically labeled metabolite species by dividing the integrated peak area of each isotope species by summed peak area of all labeled species.

**Differential expression analysis**

In order to perform RNA-seq, we back diluted six mid-log phase cultures of each strain so they would reach an OD_600_ of 0.5 after two days of growth in 10 mL of 7H9 media with antibiotic at 37° C with shaking. We pelleted each culture, removed the supernatant, resuspended the cells, and inoculated three cultures per strain with 7H9 and 50 µg/mL hygromycin or 7H12 + 0.2% w/v lactate and antibiotic. We allowed these cultures to shake at 37° C for 6 hours, then we extracted and cleaned the RNA as described above. We used the KAPA RiboErase kit (Roche, Basel, Switzerland) with Mtb custom rRNA targeting oligos for rRNA depletion. To prepare the RNA sequencing libraries, we followed the manufacturer’s instructions of the KAPA RNA HyperPrep kit (Roche). We used 25 ng of input RNA. One 7H9 -18 T>G sample was lost during the process. The prepared RNA libraries were sequenced with the MiSeq Reagent Kit v3 (150-cycle, Illumina, San Diego, CA) with 75bp paired-ended setup. Sequencing reads were mapped to the H37Rv reference genome with the bwa mem pipeline (12,14). We used the ht-seq tool ‘count’ to assign genomic features to the aligned reads; reads that aligned to rRNA were removed (12,15). To perform the differential expression analysis, we used DESeq2 according to the standard parameters (16).
